# Dopachrome tautomerase variants in patients with oculocutaneous albinism

**DOI:** 10.1101/2020.06.26.171223

**Authors:** Perrine Pennamen, Angèle Tingaud-Sequeira, Iveta Gazova, Margaret Keighren, Lisa McKie, Sandrine Marlin, Souad Gherbi Halem, Josseline Kaplan, Cédric Delevoye, Didier Lacombe, Claudio Plaisant, Vincent Michaud, Eulalie Lasseaux, Sophie Javerzat, Ian Jackson, Benoit Arveiler

## Abstract

**Purpose:** Albinism is a clinically and genetically heterogeneous condition. Despite analysis of the nineteen known genes, ∼30% patients remain unsolved. We aimed to identify new genes involved in albinism.

**Methods:** We sequenced a panel of genes with known or predicted involvement in melanogenesis in 230 unsolved albinism patients.

**Results:** We identified variants in the *Dopachrome tautomerase* (*DCT*) gene in two patients. One was compound heterozygous for a 14 bp deletion in exon 9 and c.118T>A p.(Cys40Ser). The second was homozygous for c.183C>G p.(Cys61Trp). Both patients had mild hair and skin hypopigmentation, and classical ocular features. CRISPR/Cas9 was used in C57BL/6J mice to create mutations identical to the missense mutations carried by the patients, along with one loss-of-function indel mutation. When bred to homozygosity the three mutations revealed hypopigmentation of the coat, milder for Cys40Ser compared to Cys61Trp or the frameshift mutation. Histological analysis identified significant hypopigmentation of the retinal pigmented epithelium (RPE) indicating that defective RPE melanogenesis could be associated with eye and vision defects. *DCT* loss of function in zebrafish embryos elicited hypopigmentation both in melanocytes and RPE cells.

**Conclusions:** *DCT* is the gene for a new type of oculocutaneous albinism that we propose to name OCA8.

## INTRODUCTION

In humans, albinism defines a group of genetic diseases commonly subdivided into three main types: oculocutaneous albinism (OCA), ocular albinism (OA) and syndromic albinism (Hermansky-Pudlak syndrome, HPS, and Chediak-Higashi syndrome, CHS)^1,2^. Generalized hypopigmentation of the skin and hair is highly variable among patients, ranging from total absence of melanin to undetectable defects in pigmentation. On the other hand, ophthalmological anomalies are more consistently detected and are therefore considered to be central in the definition of human albinism. These include iris and/or retinal hypopigmentation, nystagmus, foveal hypoplasia, chiasm misrouting of the optic nerves, and overall reduced visual acuity. Krujit et al. described major and minor criteria and proposed that patients should display three major criteria or two major criteria and two minor criteria for the diagnosis of albinism to be ascertained ^3^.

Albinism is also genetically heterogeneous since 19 genes are so far known to be involved (6 for OCA, 1 for OA, 1 for the closely related FHONDA, 10 for HPS, 1 for CHS) ^2,4,5^. Despite a systematic search for both single nucleotide variants and copy number variants in these 19 genes, 27% of patients remain without a molecular diagnosis ^5^.

We and others have shown that over 50% of OCA patients with molecular diagnosis have a mutation in *TYR* (OCA1) encoding tyrosinase while around 2-3% have a defective *TYRP1* (OCA3) encoding tyrosinase related 1 protein ^1,5–8^. Both enzymes are well known to control pigment synthesis ^9^. Of note, a third enzyme, dopachrome tautomerase, *DCT* acts downstream of TYR but upstream of TYRP1 on the DHICA melanin synthesis pathway ^10,11^ making the corresponding gene (*DCT/TYRP2*) an obvious candidate for human albinism. However, pathogenic variants in *DCT/TYRP2* have not yet been reported.

In the present study, we screened a series of 230 unsolved patients for pathogenic variants among a selection of candidate genes including *DCT/TYRP2*. We describe two patients with classical ocular albinism and mild hypopigmentation of the skin, hair and eyes who both have potential pathogenic *DCT/TYRP2* genotypes. We further validate these variants by analyzing the phenotypes of genetically modified mice lines and zebrafish embryos.

This work is the first evidence that mutations in *DCT/TYRP2* can cause oculocutaneous albinism in humans. We therefore propose that DCT/TYRP2 is now associated to a new OCA subgroup, OCA type 8 (OCA8).

## MATERIALS AND METHODS

### Patients

This study was approved by the local authority (Comité de Protection des Personnes Sud Ouest et Outre Mer III). Informed consent was obtained from the patients and/or their parents in the case of minors before genetic analysis was performed. Authorization for publication was obtained from Patient 1. Patient 2 was lost from contact and authorization was not collected, but the patient cannot be identified on the basis of data published here.

### Sequencing of candidate genes

DNA was extracted from peripheral blood leukocytes using an automated procedure (Tecan EVO-ReliaPrep, Promega). Next Generation Sequencing (NGS) was performed using the ionTorrent technology on a S5XL instrument (Life Technologies, ThermoFischer Scientific, U.S.A.) (AmpliSeq panel) and bioinformatics analysis of variants were as previously described ^5^. *DCT* variants (NM_001129889.2) were confirmed by Sanger sequencing. See Supplementary Data for more details.

### CRISPR modelling of *Dct* variants in the mouse

#### Mouse lines

All mouse work was carried out in compliance with United Kingdom Home Office regulations under a United Kingdom Home Office project license. To create the *Dct*118 A/A, *Dct*183 G/G and *Dct* −/− alleles the CRISPR design site https://chopchop.cbu.uib.no/ (in the public domain) was used to design guides with default settings (g1 for *Dct*118 A/A and g2 for *Dct*183 G/G and *Dct* −/−, Supplementary Materials). Genetic engineering of CRISPR-Cas9 lines with each of the Cys substitutions was performed as already described ^12^ and detailed in Supplementary Materials and Methods.

#### Direct spectrophotometry of Dct mice

Coat colour of live mice, aged between 1 and 2 months, was measured by direct spectrophotometry using Konica Minolta CM-2600d, as already described ^13^ and detailed in Supplementary Materials and Methods.

#### Eye histology

Mice were sacrificed by cervical dislocation and eyes enucleated and processed as detailed in Supplementary Materials and Methods.

### *dct* knock-down in *Danio rerio*

#### dct targeting

Zebrafish (*D. rerio*) were produced in accordance with the French Directive (Ministère de l’Agriculture) and in conformity with the European Communities Council Directive (2010/63/EU). The zebrafish homolog of *DCT* was identified by searching in the genome sequence database at http://www.ensembl.org/Danio_rerio taking into account the synteny with the human genome. Morpholino (MO) were designed by and obtained from GeneTools, LLC and embryo injection is detailed in Supplementary Materials and Methods.

*Melanin assay* was based on already described methods ^14–17^ and detailed in Supplementary Materials and Methods.

*Histological analysis* was performed as described in Supplementary Materials and Methods.

## RESULTS

### Identification of *DCT* variants in patients with oculocutaneous albinism

We recently described the analysis of 990 index cases with albinism and showed that a molecular diagnosis could be obtained after analysis of the 19 known albinism genes in 72% of patients ^5^. Our cohort nowadays comprises more than 1500 patients, amongst whom the proportion of patients remaining without a molecular diagnosis is in the 25-30% range. Following the hypothesis that some of these patients may have pathogenic variants in genes other than the 19 already identified, we screened the genomes of 230 undiagnosed patients for variants in candidate genes selected for their known or predicted involvement in melanogenesis, including orthologues of mouse mutants with pigmentary defects (see Supplementary Table 1).

Homozygous or compound heterozygous variants of the *Dopachrome tautomerase* (*DCT*) gene (NM_001129889.2), also called *Tyrosinase-related protein 2* (*TYRP2*), were identified in two patients.

Patient 1, a 12 years old girl born to unrelated parents of French origin, was diagnosed at the age of 3 years, and presented with moderate hypopigmentation of the skin and hair, congenital nystagmus, moderate foveal hypoplasia grade I ^18^, iris transillumination and hypopigmentation of the retina (Figure 1). Her visual acuity was 5/10 in both eyes. She harbored a single nucleotide variant in exon 1 of *DCT*, c.118T>A p.(Cys40Ser), that was inherited from her heterozygous father, and a 14 bp deletion in exon 9, c.1406_1419del p.(Phe469*), inherited from her heterozygous mother. Patient 2 (age at diagnosis not known) is a woman aged 36 at the time of consultation who was born to consanguineous parents originating from Northern Africa. She had cream-coloured skin, light brown hair, nystagmus, photophobia, iris transillumination, hypopigmentation of the retina and reduced visual acuity (4/10 and 5/10). Morphology of the fovea was not investigated. Patient 2 was homozygous for variant c.183C>G p.(Cys61Trp) in *DCT* exon 1 (Figure 1). Segregation analysis could not be completed and no further information on her phenotype could be gathered, as she was lost from contact after DNA sampling. All three variants were predicted to be deleterious by several prediction software (see Materials and Methods) taking into account parameters such as allele frequency in public genome databases, conservation, as well as effect on protein structure and physicochemical properties. All three variants are classified as likely pathogenic according to the criteria of the American College of Medical Genetics ^19^ : c.118T>A p.(Cys40Ser) (PS3, PM2, PP3, PP4), c.1406_1419del p.(Phe469*) (PM2, PM4, PP3, PP4), c.183C>G p.(Cys61Trp) (PS3, PM2, PP3, PP4).

**Figure 1:**
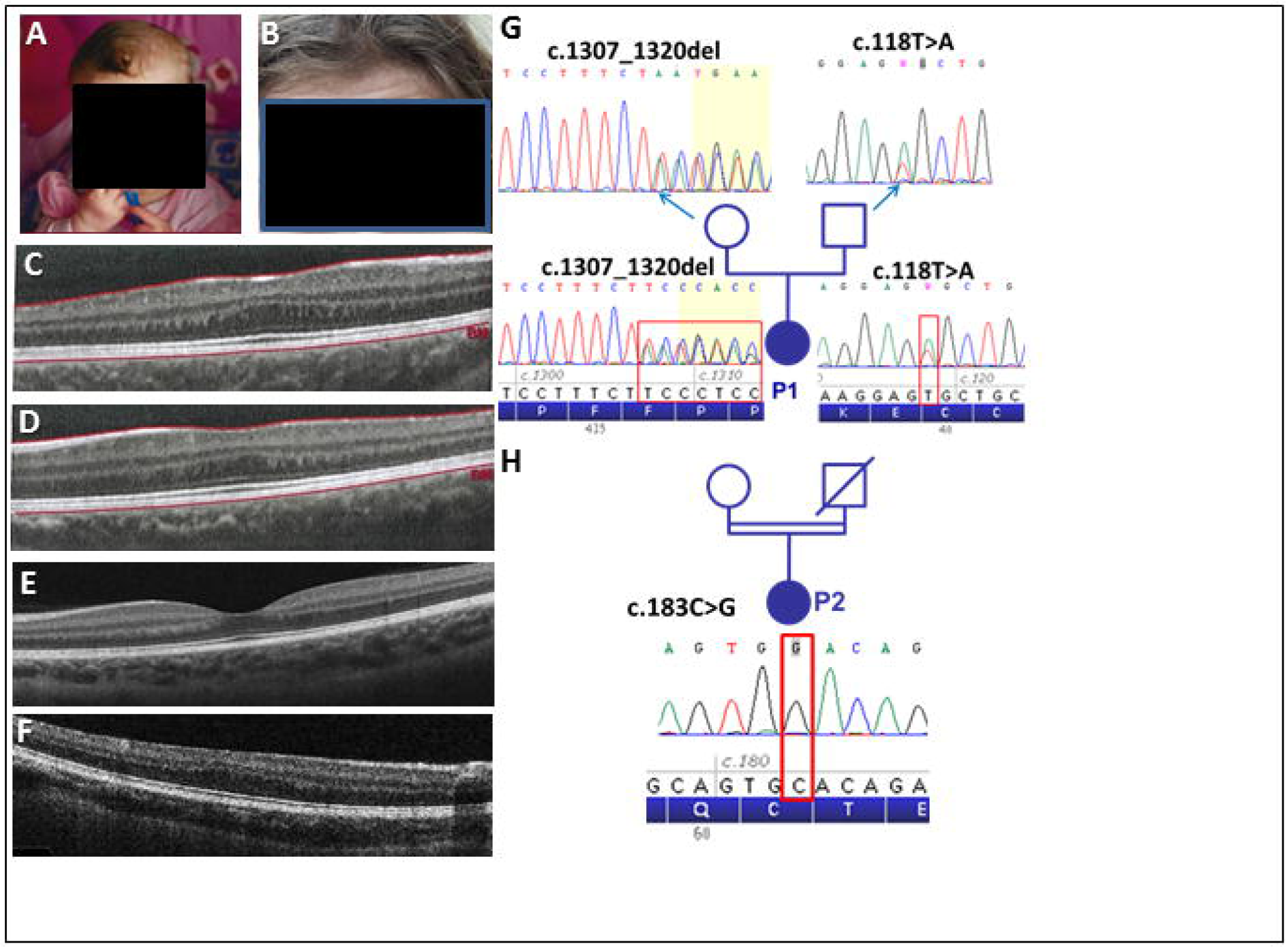
Phenotype and genotype of Patients 1 and 2. Photographs show the hypopigmentation of hair and skin of Patient 1 (A,B). Bilateral foveal hypoplasia is seen in Patient 1 (C: right eye; D: left eye), compared to a normal control (E) and a positive control (OCA1 patient) (F). Pedigree of Patient 1 shows inheritance of the two mutations from heterozygous parents (G). Pedigree of Patient 2 is shown in H. The patient’s consanguineous parents could not be tested because they were not available.

The c.1406_1419del p.(Phe469*) out of frame deletion present in Patient 1 is predicted to result either in a truncated *DCT* protein lacking the transmembrane domain which is necessary for anchoring the protein in the melanosomal membrane or in mRNA nonsense mediated decay.

Notably, both single nucleotide variants found in the two patients, c.118T>A p.(Cys40Ser) and c.183C>G p.(Cys61Trp) affect highly conserved cysteine residues located in exon 1 of the gene and belonging to a Cys-rich domain that is part of an EGF-like domain with 5 potential disulfide bonds (Figure 2).

**Figure 2.**
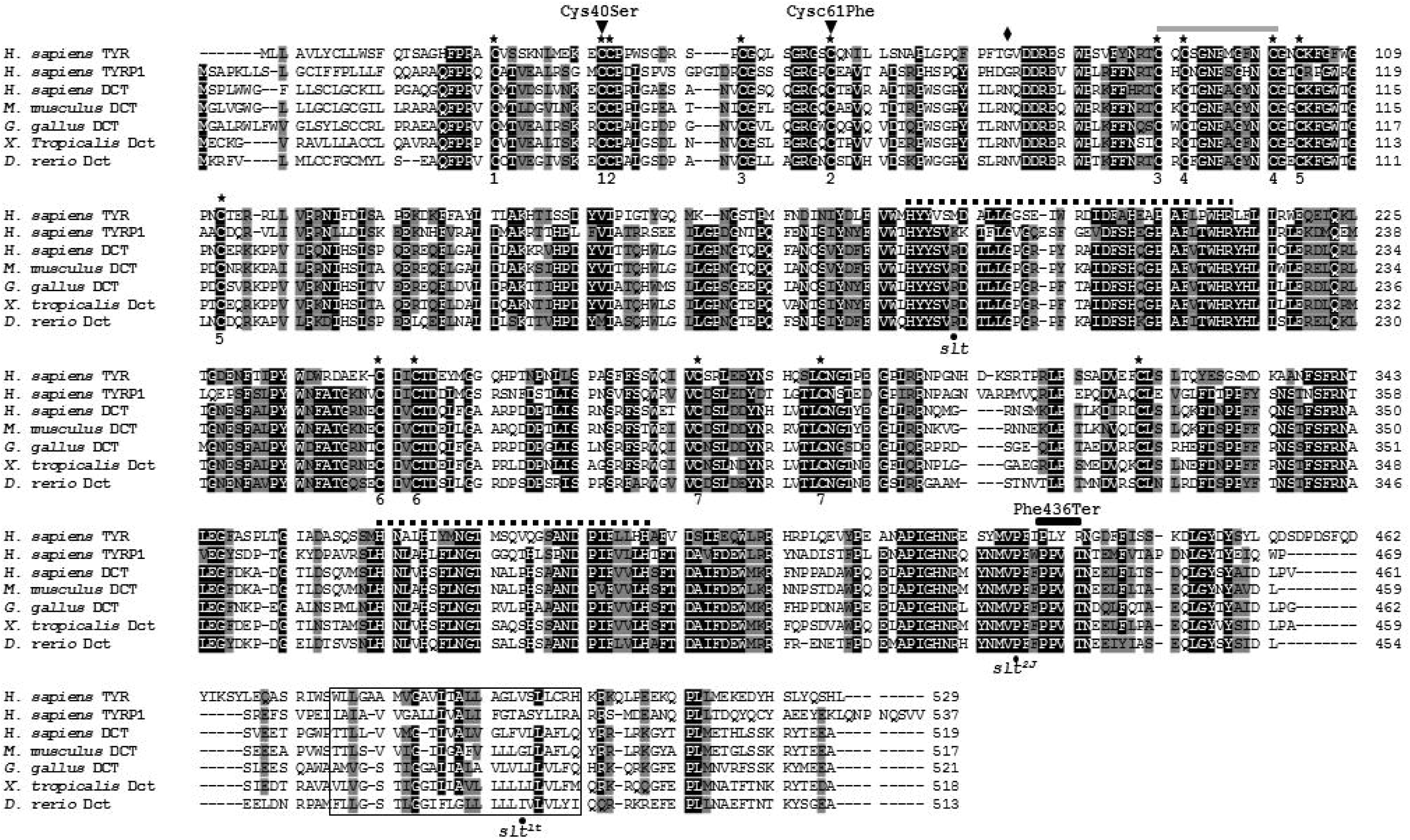
Amino acid sequences alignment for the Human tyrosinase family and DCT sequences of other species. In addition to *Homo sapiens* protein sequences of tyrosinase family including TYR, TYRP1 and TYRP2 (*DCT*), *DCT* protein sequences of *Mus musculus, Gallus gallus, Xenopus tropicalis* and *Danio rerio* were aligned. Black shaded residues indicate more than 80% of identity whereas grey shaded residues indicate more than 80% of similarity. Black stars (**\***) highlight conserved cysteine residues. Black diamond indicates the position of the fortuitous out of frame 5bp deletion in the knockout mouse model produced in the present study (p.Asn78*). Above the sequences alignment, the grey line indicates the EGF-like domain whereas the black lines indicate the metal ion-binding domains. Black dots under the alignment indicate the previously reported *slaty* mutations in mouse. Numbering under alignment indicates the cysteine bounds.

These findings suggest that the three *DCT* variants identified are pathogenic and responsible for the albinism phenotype of the two patients described here. In order to further evaluate their pathogenicity, the two missense variants were introduced in mouse by CRISP-R/Cas9 engineering.

### CRISPR modelling of *Dct* variants in the mouse

In addition to the *Dct* mouse knock-out line that was described by Guyonneau et al. ^20^, three different missense variants have been shown to be responsible for the greyish (slaty) color of mouse hair ^11,21^. As indicated in Figure 2, none of these mutations are equivalent to the patients’ variants identified in this study. We therefore decided to genetically engineer CRISPR-Cas9 lines with each of the Cys substitutions.

Two lines were obtained with the exact equivalent changes to human p.(Cys40Ser) [mouse c.118T>A p.(Cys40Ser)] and human p.(Cys61Trp) [mouse c.183C>G p.(Cys61Trp)]. Targeting of base pair 183 also led to a fortuitous out of frame 5bp deletion (c.171_175del) in one founder that was brought to homozygosity as a loss of function control *Dct−/−* (frameshift results in premature stop codon at position 78, see Supplementary data).

Mice heterozygous for any of the three mutations had a normally pigmented (black, nonagouti) fur. We crossed each line to produce homozygous mice which were grown to adulthood and all mice displayed dark-grey hair, apparently identical to that seen in the previously described mutations of *Dct* (Figure 3A). In order to more precisely quantify the pigmentary changes we colorimetrically assayed the dorsal coat of the mutants in three-dimensional CIELAB colour space, measuring along light-dark, blue-yellow and red-green axes (Figure 3B). All three lines were very different from the parental C57BL6/J mice. In addition, the Cys61Trp mice and the deletion mice were indistinguishable in all three dimensions. Interestingly, although not apparently different by eye, the mice homozygous for Cys40Ser were consistently less severely affected than the Cys61Trp and deletion mice in all three dimensions.

**Figure 3.**
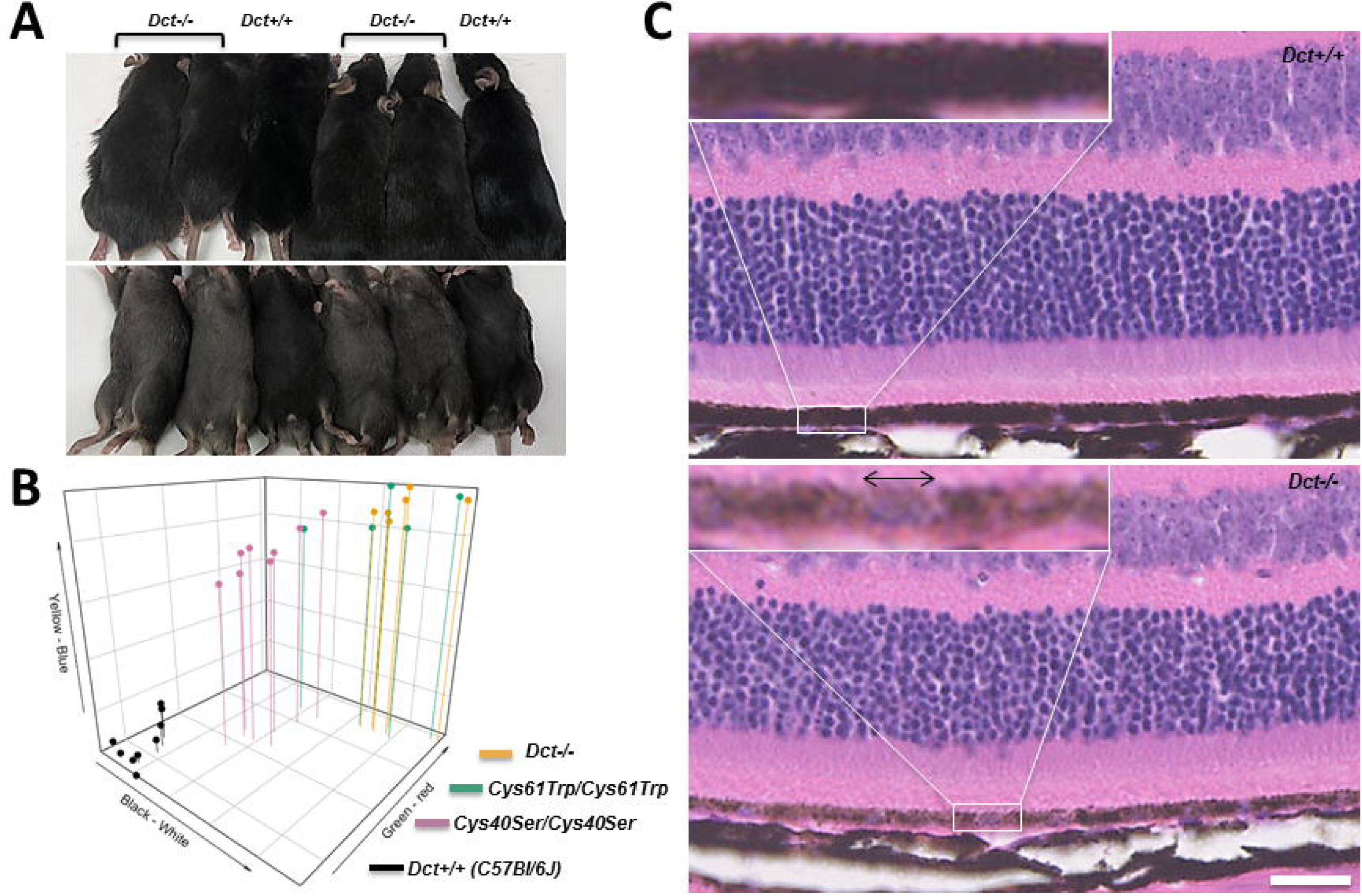
Patient mutations have a visible effect on pigmentation of the coat and eyes in mice. (A) Dorsal (upper panel) and ventral (lower panel) view of Dct−/− mice compared to C57Bl/6J controls, (B) three-dimensional CIELAB colour measurement along light-dark, blue-yellow and red-green axes as indicated for Cys40Ser, Cys61Trp, Dct−/− and C57Bl/6J. For each measurement, the difference between any of the mutated strain and control was highly significant (T-test, p-values< 10^−6^), no significant difference was found between Cys61Trp and Dct−/− whereas Cys40Ser was significantly less affected than Cys61Trp (especially in the red-green pigment contents, p-value < 5.10^−4^) (C): histology of 2 month old retinas from Dct-/*-* and age-matched C57Bl/6J control mice. Fields were photographed at mid-distance between the optic nerve and ciliary body for both samples, scale bar 25μm. One field is magnified to illustrate the significant difference in RPE pigmentation. Haematoxylin stained nuclei are clearly visible in Dct−/− (double arrow line) whereas nuclei are masked by intense pigmentation in control.

These observations in mice strongly suggest that the missense variants identified in both patients cause loss or partial loss of function of *DCT* and that this is the cause of their hypopigmentation of the hair and skin.

*Dct* knock-out and *slaty* mice have hardly been investigated for ophthalmological anomalies. We conducted histological analysis of eyes from adult *Dct* −/− mice. Gross anatomy of the eye was not modified in these mice as compared to wild-type (C57Bl/6J) age-matched controls although we cannot rule out that the outer nuclear layer was slightly thinner and disorganized in some *Dct−/−* retinas (not shown). Melanocytes in the choroidal layer appeared normally pigmented. However, careful examination of the outer retina evidenced that the RPE in *Dct−/−* mice was significantly less pigmented than in wild-type retinas. As illustrated in Figure 3C, dark melanin in the control eye completely masks the RPE cell membranes and contents whereas in *Dct−/−* eyes, pigmentation of the RPE is light enough that hematoxylin stained nuclei can be visualized in most of the RPE cells.

Altogether, these results show that when introduced in mice, Cys40Ser and Cys61Trp mutations result in hypopigmentation of the coat, the latter being undistinguishable from complete loss of function. Furthermore, RPE melanin content is reduced in *Dct−/−* mice, which may cause eye and vision anomalies in mice, and, in a similar way, in *DCT* patients.

### *dct* knock-down in *Danio rerio*

Loss of function of *dct* was also investigated in zebrafish (*D. rerio*) embryos in which development and pigmentation of skin melanophores and RPE can be followed *in vivo*. Of note, only one *DCT* homologue is found in *D. rerio* genome and *D. rerio* protein sequences are highly conserved with 62% identity and more than 85% similarity (Figure 2). The expression of *dct* was transiently knocked down using a morpholino (MO). Compared with embryos injected with MO-ctrl, MO-*dct* injected embryos did not show macroscopic phenotype disruption and no obvious disruption of skin melanophores pattern at neither 48 hpf nor 120 hpf (Figure 4A,C). However, melanophores dorsally localized at the head level of 48 hpf MO-*dct* appeared lighter and wider than those of controls (Figure 4A). Eye pigmentation was also slightly lighter in *dct* knocked-down embryos (Figure 4A). In order to quantify a visible hypopigmentation, a whole-embryo melanin assay was performed, confirming a ≈30% significant decrease of total melanin in *dct*-knockdown embryos versus control embryos (Figure 4B). Later in development at 120 hpf, MO-*dct* embryos still presented with a normal pattern of skin melanophores. However, their intracellular pigmentation was different from controls. In control embryos, pigmentation of melanophores appeared highly packed as expected whereas pigmentation in MO-*dct* embryos melanophores was consistently weaker and showed an abnormal stellar pattern (Figure 4C). In addition to general pigmentation disruption, significant microphthalmia was noticed possibly due to off-target effect or toxicity, although other indicators for toxicity ie shape and size of the embryos, were not significantly affected^22^ (Figure 4D). Histological sectioning of 48 hpf embryos (Figure 4A) evidenced eye developmental defects. In MO-Ctrl injected embryos, stratification of the neural retina into outer and inner layers was achieved by 48 hpf whereas at the same stage, MO-*dct* eyes still displayed only one homogeneous layer of neurons. The RPE in morphants was clearly hypopigmented, but in addition instead of the normal organization as a heavily pigmented cobblestoned single cell layer, the RPE showed dishevelled cells and wavy contours.

**Figure 4.**
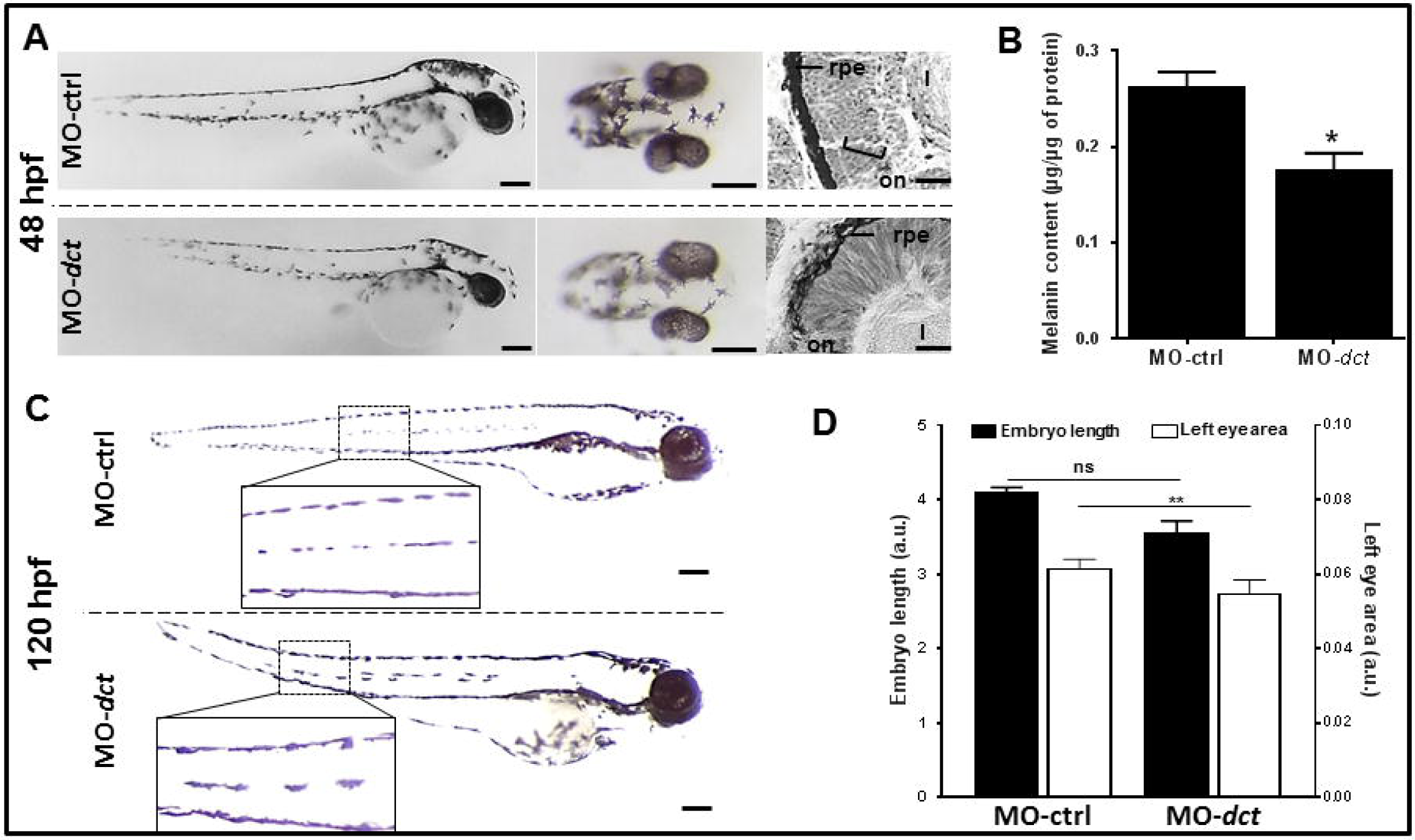
*dct* is required for skin melanocytes and RPE development preservation in zebrafish *D. rerio*. (A) Representative lateral views of whole embryos, dorsal views of heads and histological cross sections of eyes from representative 48hpf zebrafish embryos injected with MO-ctrl or MO-*dct*. (B) Melanin was quantified in 48hpf zebrafish embryos injected with MO-ctrl or MO-*dct*. Means were calculated from the data collected from three independent experiments, Mann-Whitney test, *p < 0.05. (C) Representative lateral views of whole embryos at 120 hpf injected with MO-ctrl or MO-*dct*. Insets represent enlarged views of skin melanocytes laterally localized in the trunk. (D) Embryo length (reported on left y axis) and left eye area (right y axis) were measured from one representative experiment on MO-ctrl group (n=12) and MO-*dct* group (n=28). Mann-Whitney test was applied for length data whereas unpaired t-test was performed for left area data, ns, non-significant, **p<0.01.

Altogether, these data are complementary to the observations made in mice, and confirm that loss of function of *Dct* impairs melanin metabolism in both melanocytes and in RPE cells of early embryos in *D. rerio* with possible consequences on eye development and/or function.

## DISCUSSION

We report the identification of pathogenic variants in the *DCT* gene in two unrelated patients with oculocutaneous albinism. Two of them are missense variants affecting highly conserved cysteine residues that belong to a Cys-rich domain [c.118T>A p.(Cys40Ser) and c.183C>G p.(Cys61Trp)]. The third variant is a 14 bp out-of-frame deletion [c.1406_1419del p.(Phe469*)], a likely null allele. Both patients presented with a creamy skin and light brown hair, thus indicating that melanin is produced to a significant level. Visual acuity was moderately reduced (4/10 – 5/10) in both patients. The ocular phenotype included nystagmus, hypopigmentation of the retina and iris transilllumination. Foveal hypoplasia was of low grade in patient 1 and not investigated in patient 2.

This is the first time *DCT* variants have been shown to cause oculocutaneous albinism in human. Dopachrome Tautomerase (*DCT*), formerly known as Tyrosinase Related Protein 2, is one of three members of the tyrosinase protein family, along with TYR and TYRP1. All three are enzymes catalysing reactions in the melanogenesis pathway, and are located within melanosomes, a pigment cell-specific organelle ^9^. In this series of reactions as previously reviewed^9^, Tyrosinase (TYR) catalyses the initial conversion of tyrosine to dihydroxyphenylalanine (DOPA) and DOPA to DOPAquinone, which spontaneously cyclises to DOPAchrome. *DCT* catalyses the tautomerisation of DOPAchrome to dihydroxyindole carboxylic acid (DHICA). DOPAchrome can also spontaneously lose the carboxyl group to produce dihydroxyindole (DHI). The function of the third member of the family, Tyrosinase-related Protein 1 (TYRP1) is still not completely certain and may vary depending upon species and/or cell types. It probably acts, maybe in conjunction with TYR, to oxidise DHICA and DHI, by removing the two hydroxyl groups to the indole derivatives which ultimately polymerise to form melanin on a protein matrix within the melanosome. The melanins formed from DHICA and DHI are eumelanins, which range in colour from brown to black.

As TYR is the key limiting enzyme in melanogenesis, its absence results in complete lack of melanin, as seen in mutant mice and humans with OCA1. Absence of TYRP1 in mice results in the production of brown eumelanin, which appears to be less polymerised than black eumelanin. The human *TYRP1* gene is mutated in patients with OCA3. The effect on pigmentation, and ocular defects, are relatively mild with bright copper-red coloration of the skin and hair, both in African and European patients ^5,23^. Curiously, despite being upstream of TYRP1, the absence of *DCT* in mice results in a darker eumelanin than the brown eumelanin that lacks end-products generated in the presence of TYRP1. This is most likely due to DOPAchrome being converted to DHI, which produces a dark eumelanin. Absence of TYRP1 in presence of *DCT* means that DHICA, rather than its oxidised derivative, is polymerised to produce the brown eumelanin observed. A milder phenotype than the average OCA3 type is therefore expected in humans with a loss of *DCT* function.

The two missense mutations identified in this study are located in the N terminal Cys-rich subdomain of *DCT* which is strikingly conserved in its two paralogues TYR and TYRP1 and across species (Figure 2). Localisation of the ten cysteine residues (C1 to C10) in each enzyme is indicated in Table 1 as well as their participation in the 5 disulfide bonds that stabilize the domain ^9^.

**Table 1:**
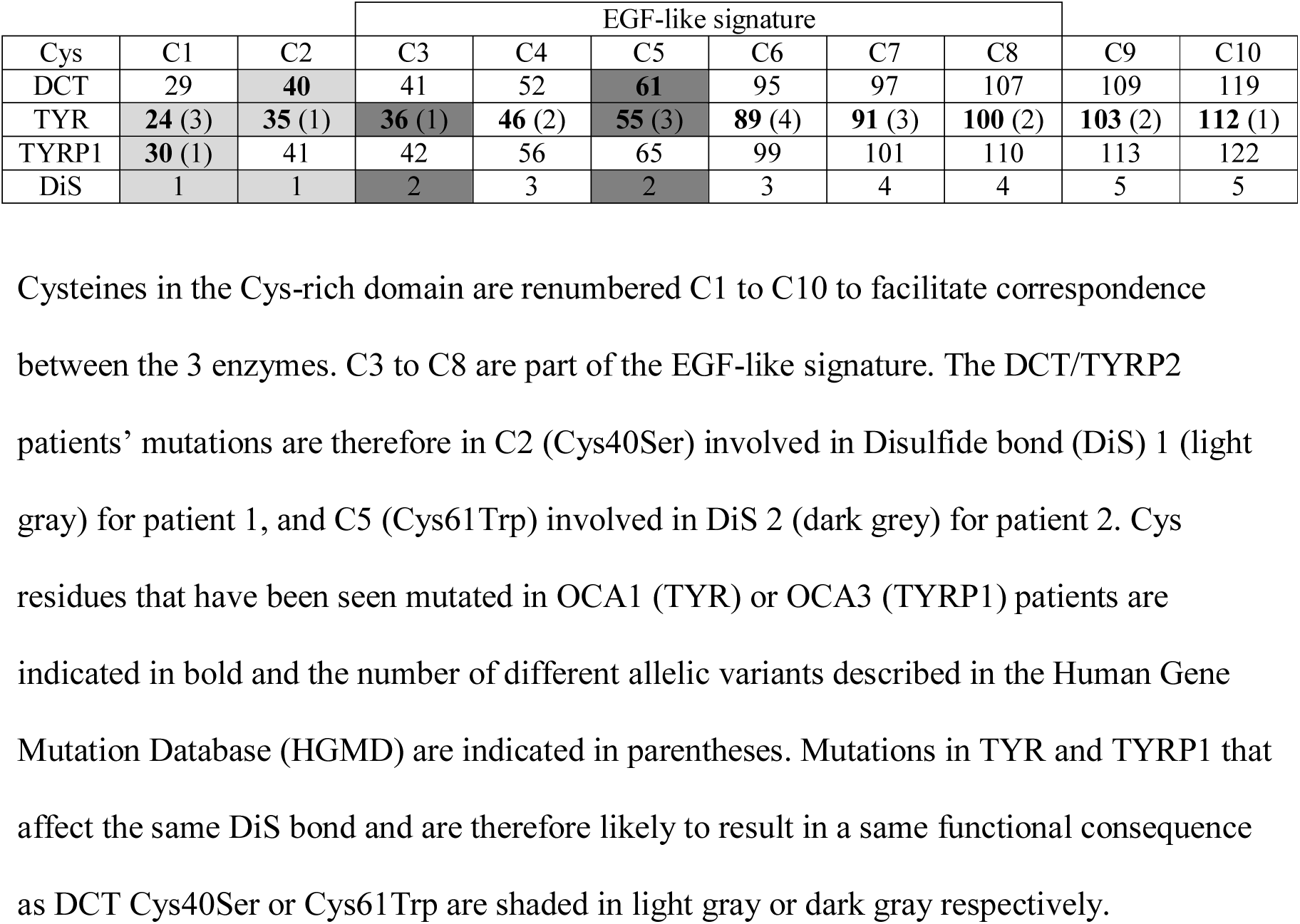
Conserved Cysteine residues across the TYR/TYRP1/*DCT* family.

Note that C3 to C8 (involved in disulfide bonds 2, 3, 4) belong to the highly conserved EGF-like signature. According to the Human Gene Mutation Database, amongst 354 missense and nonsense mutations in TYR, substitutions in all of the C1 to C10 cysteine residues are found associated with OCA1. Of particular interest, C2 [patient 1 variant p.(Cys40Ser)] is mutated in two unrelated OCA1 Pakistani patients ^24^, and C5 [patient 2 variant p.(Cys61Trp)] is also mutated in OCA1 patients with 3 identified variants ^25–27^. Interestingly one of the OCA1 patients with a C2 substitution described by Jaworek et al. is homozygote for a Cys to Arg change and has a somewhat mild phenotype (fair blond hair and pinkish skin) compared with known complete loss of function. Moreover, this mutated TYR has a temperature-sensitive behaviour likely due to slight protein misfolding. Whether this is due to loss of disulfide bond 1 and is therefore to be expected whatever amino acid is substituted at C1 or C2 would require further structural or functional investigations. Interestingly, the only allele involving a cysteine change of the Cys-rich domain in OCA3 patients affects C1 of TYRP1 and has also been associated with impairment of traffic to the melanosomes ^28^.

Altogether, these data suggest that disruption of disulfide bond 1 in TYR and/or TYRP1 is likely to result in less severe and possibly temperature sensitive misfolding of the mutated enzyme. Ablation of bond 2 on the other hand may have a more detrimental effect on the structure of the whole enzyme as it is essential to stabilize the EGF-like domain.

Given the high degree of conservation of tyrosinase related proteins between them and among species, we speculated that substitution of C2 and C5 in our *DCT* patients could have similar effects and investigated the corresponding phenotypes in mice. Supporting the relevance of this model for our comparative functional analysis, the two classical mutations of mouse in *Tyr* (albino) and *Tyrp1* (brown) are also in conserved cysteines of the Cys-rich domain, in tyrosinase at C9, Cys103Ser, ^29,30^ and in Tyrp1 at C8, Cys110Tyr ^31^. We show that mutation of *Dct* C2 in mice appears to be less severe than either C5 or the null frameshift that was produced. Although the difference is not readily detectable visually, the subtle variation is clearly seen when the fur is analysed spectrophotometrically. It is tempting to attribute these differences to only a slight conformation change as in TYR and TYRP1. However structural models of *DCT*/TYRP2 are speculative at this point as no crystallographic data have been so far generated.

Altogether these observations indicate a causal role of the C2 and C5 substitutions in our patients. The mild phenotype of patient 1 may be explained by a partial and possibly thermosensitive loss of function of *DCT*.

Previous studies ^20^ have reported no effect of *Dct* mutations on mouse eye pigmentation. However, our close examination of mice with the null frameshift reveals a significant decrease in pigmentation of the RPE, although the choroid appears only mildly or not affected at all. How can this be related to some or all of the ophthalmic anomalies seen in the patients? Since our original demonstration that the RPE plays a key role in early development of the mammalian neural retina ^32^, a considerable amount of work has been performed, including in the albino *Tyr−/−* mouse, that sheds light on the primary cause for visual deficiency in OCA1 patients [for review see ^33^]. In particular it was shown that the lack of melanin and/or intermediates along the melanogenesis pathway leads to retinal developmental defects, especially in the neurogenesis of ganglion cells, and that defective guidance of their axons along the optic tracts results in optic chiasm misrouting and impairment of binocular vision [for review see ^34^]. Despite extensive investigation using partial rescue of tyrosinase activity in the *Tyr−/−* model ^35–38^, it is still not clear which factors in the pigment synthesis pathway are involved and if they directly signal from the RPE to the neural retina or if they primarily affect RPE integrity. The fact that the RPE of both *Dct−/−* mice and zebrafish *dct* morphants is hypopigmented, together with typical ophthalmological defects diagnosed in our two DCT patients supports the hypothesis that the fully functional melanin pathway in the early RPE is instrumental to retinal neurogenesis.

The zebrafish embryo is a useful experimental model to study both skin and eye pigmentation as well as early eye development. Models for OCA1 and OCA3 have been described ^39–45^. Although *dct* expression pattern in zebrafish embryo has been used as a molecular marker of pigmented cells ^44,46^, its knock-down has not been previously described. Here we show that it results in hypopigmentation of both melanophores and RPE. In addition, instead of harbouring their usual pigmentation pattern, melanophores of the skin resemble those of wild-type embryos responding to hormone-driven adaptation to surrounding darkness by melanin dispersion ^41^, suggesting that *dct* knock-down may impair melanin aggregation or that MO treatment has affected intracellular signalling. In the RPE, in addition to hypopigmentation, cell integrity was obviously affected. Such a pleiotropic effect has been described previously ^40^. Further investigations will aim at confirming and studying these changes at the cellular and molecular levels during development both in mouse and zebrafish embryos.

In conclusion, we report here the first evidence of *DCT* variants causing OCA in two unrelated patients. It is notable that both patients had mild hypopigmentation of the skin, hair and eyes and moderate loss of visual acuity. Albinism is clinically heterogeneous, and *DCT* patients seem to fall in the mild range of the phenotypical spectrum. These mild phenotypes may be overlooked by clinicians and lead to underdiagnosis of albinism. As we recently discussed ^47^, we advocate that minimal signs, both dermatologic and ophthalmologic, sometimes of atypical presentation, should be systematically given consideration and lead to genetic investigations in order to establish a precise diagnosis. The *DCT* gene should now be included in albinism diagnosis gene panels. We propose that the corresponding form of OCA is called OCA8.

## ACKNOWLEDGEMENTS

The authors are grateful to the French Albinism Association Genespoir for financial support and for timeless action in favour of patients with albinism. IG, LM, MK and IJJ were funded by the MRC University Unit award to the MRC Human Genetics Unit, grant number MC_UU_00007/4. They also wish to thank the patients for participating in this study and Angus Reid for assisting with colorimetric measurements.

## AUTHORSHIP CONTRIBUTIONS

Perrine Pennamen: Conceptualization, Next Generation Sequencing, Bioinformatics, Writing – original draft

Angèle Tingaud-Sequeira: Zebrafish experiments

Iveta Gazova: CRISPR/Cas9 editing, coat spectrophotometric analysis

Margaret Keighren: CRISPR/Cas9 editing, coat spectrophotometric analysis

Lisa McKie: Mouse eye histology

Sandrine Marlin: Patients investigations

Gherbi Halem: Patients investigations

Josseline Kaplan: Patients investigations

Cédric Delevoye: Conceptualization

Claudio Plaisant: Next Generation Sequencing

Vincent Michaud: Patients cohort characterization

Didier Lacombe: Supervision

Eulalie Lasseaux: Patients cohort characterization

Sophie Javerzat: Conceptualization, Eye histology interpretation (mouse and zebrafish), Visualization, Writing – original draft, review and editing

Ian Jackson: Conceptualization, Visualization, Writing – original draft, review and editing, Funding acquisition

Benoit Arveiler: Conceptualization, Project administration, Supervision, Visualization, Writing – original draft, review and editing, Funding acquisition

