## Supplementary Materials and Methods for "Dopachrome tautomerase variants in patients with oculocutaneous albinism"

**Sequencing of the candidate genes, bioinformatics analysis.**

DNA samples were quantified using the Qubit™ dsDNA HS Assay Kit on the Qubit® 2.0 Fluorometer (Life Technologies) and diluted to a final concentration of 10 ng/µL. Amplicons (libraries) were generated with the Ion AmpliSeq™ Library Kit 2.0 and then quality-checked and quantified on the 2200 TapeStation (Agilent, U.S.A.). Emulsion PCR and enrichment were performed on the Ion CHEF System (Life Technologies) using the Ion 520/530 Kit Chef. Two Ion 530 chips (Life Technologies), each containing 24 patients, were sequenced per run.

Pathogenicity of single nucleotide variants was evaluated using bioinformatics tools Cartagenia BenchLab NGS (Agilent Technologies) and Alamut Visual (Interactive BioSoftware, Rouen, France) based on criteria including allele frequency (gnomAD http://gnomad.broadinstitute.org, ESP evs.gs.washington.edu/EVS, dbSNP www.ncbi.nlm.nih.gov/SNP), description in database (HGMD Professional www.biobase-international.com/product/hgmd), conservation, pathogenicity prediction algorithms (Polyphen genetics.bwh.harvard.edu/pph2, SIFT sift.bii.a-star.edu.sg, Mutation Taster www.mutationtaster.org, GVGD agvgd.iarc.fr) and effect on splicing (SSF www.genet.sickkids.on.ca/~ali/splicesitefinder.html, HSF www.umd.be/HSF3, MaxEntScan genes.mit.edu/burgelab/maxent/Xmaxentscan_scoreseq.htm, NNSplice www.fruitfly.org/seq_tools/splice.html, GeneSplicer ccb.jhu.edu/software/genesplicer).

*DCT* variants (NM_001129889.2) found by NGS were confirmed by Sanger sequencing (BigDye® Terminator v3.1 Cycle Sequencing Kit; Applied Biosystems®, ThermoFischer Scientific) on a 3500xL Dx Sequencer (Life Technologies, ThermoFischer Scientific) using primers AGGAGAAAAGTACGACAGAGACA and TCCGACCAAAACCATCATTG (exon 1) and ACAGTCAGGTTGGTGTGATC and TCAGCTCATTAGTCCCTTTCTGA (exon 9).

**CRISPR modelling of *Dct* variants in the mouse.**

*Mouse lines:* To create the *Dct*118 A/A, *Dct*183 G/G and *Dct* -/- alleles the CRISPR design site https://chopchop.cbu.uib.no/ (in the public domain) was used to design guides with default settings (g1 for *Dct*118 A/A : CTTGGATGGCGTGCTGAACAAGG ; g2 for *Dct*183 G/G and *Dct* -/- : GTGGATTTCTAGAGGGCAGGGGG). Guides were *in vitro* transcribed using GeneArt Precision gRNA kit (Invitrogen, A29377). Guide RNAs along with GeneArt Platinum Cas9 Nuclease (Invitrogen, B25641) and repair single stranded oligo DCT118 or DCT183 (DCT118: GGTTGTCTGGGCTGCGGAATTCTGCTCAGAGCTCGGGCTCAGTTTCCCCGAGTCTGCATGACCTTGGATGGCGTGCTGAACAAAGAAAGCTGCCCGCCTCTGGGTCCCGAGGCAACCAACATCTGTGGATTTCTAGAGGGCAGGGGGCAGTGCGCAGAGGTGCAAACAGACACCAGACCCTGGAGTGGCCCTTATATCCT; DCT183: CAGTTTCCCCGAGTCTGCATGACCTTGGATGGCGTGCTGAACAAGGAATGCTGCCCGCCTCTGGGTCCCGAGGCAACCAACATCTGTGGATTTCTAGAGGGCAGGGGACAGTGGGCAGAGGTGCAAACAGACACCAGACCCTGGAGTGGCCCTTATATCCTTCGAAACCAGGATGACCGTGAGCAATGGCCGAGAAAATT, synthetized by IDT), was used for pronuclear injection into C57BL/6J zygotes. The injected eggs were cultured overnight to the two-cell stage and then transferred into pseudopregnant females. Pups were screened for the mutations by sequencing PCR fragments generated using primers Dct1_F and Dct1_R (Dct_1_F GCCAGGCCTCCCAATTAAGA and Dct_1_R GCCCTCACCTGTGCATTTG) and lines established carrying the targeted missense mutations and one 5 nucleotide deletion, resulting in frameshift. Genotyping was done by PCR, and then Sanger sequencing, using primers Dct1_F and Dct1_R. All lines were maintained on the C57BL/6J mouse strain background.

*Direct spectrophotometry of Dct mice:* Coat colour of live mice, aged between 1 and 2 months, was measured by direct spectrophotometry using Konica Minolta CM-2600d. Measurement of mouse’s dorsal coat colour was carried out in triplicate and the mean values of these measurements were plotted onto a 3D graph of CIELAB colour space using Rstudio with plot3D package.

*Eye histology:* Mice were sacrificed by cervical dislocation and eyes enucleated and placed into Davidson's fixative (28.5% ethanol, 2.2% neutral buffered formalin, 11% glacial acetic acid) for overnight. Prior to wax embedding, eyes were dehydrated through an ethanol series. Hematoxylin and eosin staining was performed on 5- or 10-μm paraffin-embedded tissue sections and images captured using a Nanozoomer XR scanner (Hamamatsu, Hamamatsu City, Japan) and viewed using NDP.view2 software (Hamamatsu).

***dct* knock-down in *Danio rerio*.**

*dct targeting:* Zebrafish (*D. rerio*) were produced in our facilities, in accordance with the French Directive (Ministère de l'Agriculture) and in conformity with the European Communities Council Directive (2010/63/EU). The zebrafish homolog of human *DCT* was identified by searching in the genome sequence database at http://www.ensembl.org/Danio_rerio taking in account the synteny with the human genome. Morpholinos (MO) were designed by and obtained from GeneTools, LLC. A specific MO to *dct* gene, MO-DCT, was designed as complementary to the sequence flanking the translation initiation codon of *dct* (5’-GCGCTT[CAT]TTCTGCGAAAACACAG-3’, brackets indicate the initiation codon complementary sequence). Standard control MO from Gene Tools LLC (5’-CCTCTTACCTCAGTTACAATTTATA-3’) was used as negative control. MOs were injected at the 1-2 cells stage with 1.5ng/embryo. Embryos were then incubated at 28°C. Embryos were imaged under a stereomicroscope. Length and left eye area were measured using ImageJ software.

*Melanin assay:* Ten embryos at 48 hpf stage were transferred in Tris pH=7.5 50mM, EDTA pH=8 2mM, NaCl 1mM, Triton X-100 1% and incubated 5 min at 4°C for lysis. Lysates were centrifuged for 10 min, 14000 rpm, 4°C. Supernatants were processed for classical protein assay. Pellets were dissolved in 1N NaOH. Standard solutions of melanin and samples were heated at 100°C for 10 min and processed for melanin assay by absorbance determination at 490 nm. Melanin content was normalized to protein content.

*Histological analysis:* Zebrafish embryos were fixed with 4% PFA in PBS at 4 °C overnight, embedded in paraplast and sectioned at 10 µm. After rehydration, sections were stained with toluidine blue 0.2% for 1 min and mounted in Eukitt mounting medium.

**Supplementary Table 1: Candidate genes tested in this study.**

| **GENE** | **PROTEIN** | **MOUSE MUTANT** |
| --- | --- | --- |
| *AP1B1* | Adaptor Protein 1 complex subunit beta-1 |  |
| *AP1G1* | Adaptor Protein 1 complex subunit gamma-1 |  |
| *AP1G2* | Adaptor Protein 1 complex subunit gamma-2 |  |
| *AP1M1* | Adaptor Protein 1 complex subunit mu-1 |  |
| *AP1M2* | Adaptor Protein 1 complex subunit mu-2 |  |
| *AP1S2* | Adaptor Protein 1 complex subunit sigma-2 |  |
| *AP1S3* | Adaptor Protein 1 complex subunit sigma-3 |  |
| *AP3B2* | Adaptor Protein 3 complex subunit beta-2 |  |
| *AP3M1* | Adaptor Protein 3 complex subunit mu-1 |  |
| *AP3M2* | Adaptor Protein 3 complex subunit mu-2 |  |
| *AP3S1* | Adaptor Protein 3 complex subunit sigma-1 |  |
| *AP3S2* | Adaptor Protein 3 complex subunit sigma-2 |  |
| *ATP7A* | ATPase copper transporting alpha | mottled |
| *BLOC1S1* | Biogenesis of lysosome-related organelles complex subunit 1 |  |
| *BLOC1S4* | Biogenesis of lysosome-related organelles complex subunit 4 | cappuccino |
| *BLOC1S5* | Biogenesis of lysosome-related organelles complex subunit 5 | muted |
| *DCT* | Dopachrome tautomerase | slaty |
| *DOCK7* | Dedicator of cytkinesis 7 | misty |
| *FIG4* | Phosphoinositide 5-Phosphatase | pale tremor |
| *KIF13A* | Kinesin Family Member 13A |  |
| *PMEL* | Premelanosome Protein | silver |
| *RAB38* | Ras-related protein RAB38 | chocolate |
| *RAB27A* | Ras-related protein RAB27A | ashen |
| *RABGGTA* | Rab geranyl geranyl transferase | gunmetal |
| *SLC7A11* | Solute carrier family member 11 | subtle gray |
| *VPS33A* | Vacuolar protein sorting-associated protein 33A | buff |
